# Host starvation and *in hospite* degradation of algal symbionts shape the heat stress response of the *Cassiopea*-Symbiodiniaceae symbiosis

**DOI:** 10.1101/2023.06.12.544603

**Authors:** Gaëlle Toullec, Nils Rädecker, Claudia Pogoreutz, Guilhem Banc-Prandi, Stéphane Escrig, Christel Genoud, Cristina Martin Olmos, Jorge Spangenberg, Anders Meibom

## Abstract

Global warming is causing large-scale disruption of cnidarian-Symbiodiniaceae symbioses fundamental to major marine ecosystems, such as coral reefs. However, the mechanisms by which heat stress perturbs these symbiotic partnerships remain poorly understood. In this context, the upside-down jellyfish *Cassiopea* has emerged as a powerful experimental model system. We combined a controlled heat stress experiment with isotope labeling and correlative SEM–NanoSIMS imaging to show that host starvation is a central component in the chain of events that ultimately leads to the collapse of the *Cassiopea* holobiont. Heat stress caused an increase in catabolic activity and a depletion of carbon reserves in the unfed host, concurrent with a reduction in the supply of photosynthates from its algal symbionts. This state of host starvation was accompanied by pronounced *in hospite* degradation of algal symbionts, which may be a distinct feature of the heat stress response of *Cassiopea*. Interestingly, this loss of symbionts by degradation was to a large extent concealed by body shrinkage of the starving animals, resulting in what could be referred to as ’invisible’ bleaching. Overall, our study highlights the importance of the nutritional status in the heat stress response of the *Cassiopea* holobiont. Compared with other symbiotic cnidarians, the large mesoglea of *Cassiopea*, with its structural sugar and protein content, may constitute an energy reservoir capable of delaying starvation. It seems plausible that this anatomical feature at least partly contributes to the relatively high stress tolerance of these animals in our warming oceans.

## Introduction

The immense diversity and abundance of photosymbioses between cnidarian animal hosts and their intracellular dinoflagellate algal symbionts is testimony to their ecological success in the (sub-)tropical oceans of today (Davy et al., 2012; LaJeunesse et al., 2018). In these cnidarian-algal symbioses, the algal symbionts fix large amounts of inorganic carbon through photosynthesis and subsequently transfer a substantial proportion of their photosynthates to their host (Falkowski et al., 1984; Freeman et al., 2016; Kopp et al., 2015; Muscatine et al., 1981; Rädecker & Meibom, 2022; Tremblay et al., 2012). This efficient transfer of photosynthates fuels the energy metabolism of the cnidarian host and can be sufficient to cover its respiratory carbon requirements (Cunning et al., 2017b). In turn, the cnidarian host supplies the algal symbionts with inorganic nutrients from its own metabolism, CO_2_ in particular (Leggat et al., 1999; Rädecker et al., 2017). This efficient recycling of organic and inorganic nutrients enables the cnidarian-algal symbioses to thrive even in highly oligotrophic tropical environments (Muscatine & Porter, 1977).

Yet, this ecological success is threatened by global climate change, especially global warming (Hughes, Barnes, et al., 2017). Severe and prolonged heat stress can disturb this symbiosis, ultimately resulting in its breakdown (Hughes, Kerry, et al., 2017; Lesser, 1997). This susceptibility is particularly apparent in the coral-algal symbiosis that has constituted the functional basis of coral reef ecosystems over hundreds of millions of years (Falkowski et al., 1984; Frankowiak et al., 2016; LaJeunesse et al., 2018; Muscatine et al., 1981, 1989). The breakdown of this symbiosis due to heat stress, referred to as coral bleaching, results in the loss of algal symbiont cells and photosynthetic pigments, which makes the corals tissue appear transparent revealing the underlying white calcium carbonate skeleton (Jones & Hoegh-Guldberg, 1998). During prolonged heat stress, bleaching thus deprives the host of its primary energy source, the algal photosynthates, typically resulting in starvation and, ultimately, death. Indeed, repeated mass bleaching events have caused mass mortality of corals in the last decades, pushing many coral reefs to the brink of ecological collapse (Hoegh-Guldberg, 1999; Hoegh-Guldberg et al., 2007; Hughes et al., 2018; Hughes, Kerry, et al., 2017).

The breakdown of the cnidarian-algal symbiosis is not only the result of the thermal tolerance limits of either symbiotic partner but depends also on their interactions (Bhagooli & Hidaka, 2003; Ventura et al., 2018). In this context, recent studies have shown that the destabilization of symbiotic nutrient exchange precedes the breakdown of the coral-algal symbiosis during heat stress (Baker et al., 2018; Gibbin et al., 2018; Hoadley et al., 2015; Rädecker et al., 2021). It has also been established that heterotrophic feeding enhances the tolerance and/or resilience of the coral-algae symbiosis to heat stress (Anthony et al., 2009; Grottoli et al., 2006; Tremblay et al., 2016). Hence, the heat tolerance of these animals appears to be intimately linked to the nutritional status of the host. However, the role of nutrient cycling and the nutritional state of symbiotic partners in the breakdown of cnidarian-algal symbiosis during heat stress remains poorly documented and understood.

*Cassiopea,* a genus of symbiotic jellyfish (Schyphozoa, Rhizostomae), is an emerging model organism for the study of cnidarian-algal symbiosis (Lampert, 2016; Medina et al., 2021; Ohdera et al., 2018). Similar to corals, *Cassiopea* is associated with dinoflagellates of the family Symbiodiniaceae. However, in contrast to corals and sea anemones, algal symbionts in *Cassiopea* medusae are primarily hosted by specific cells called amoebocytes within the mesoglea (Colley & Trench, 1985; Lyndby et al., 2020). From within these amoebocytes, the algae provide their host with photosynthates, thereby enabling medusae anabolic growth (Cates, 1975; Freeman et al., 2016; Hofmann & Kremer, 1981; Lyndby et al., 2020; Verde & McCloskey, 1998; Welsh et al., 2009). In contrast to corals, however, *Cassiopea* appears to thrive even under rapidly changing environmental conditions and has recently been described as invasive in many (sub-)tropical regions (Cillari et al., 2022; Holland et al., 2004; Morandini et al., 2017; Thé et al., 2021). Indeed, *Cassiopea* is now known to be relatively heat tolerant, i.e., they bleach at higher temperatures than most reef-building corals (Aljbour et al., 2019; Banha et al., 2020; Béziat & Kunzmann, 2022; Klein et al., 2019; McGill & Pomory, 2008; Tilstra et al., 2022). In part because of this relatively high heat tolerance and a high trophic plasticity, *Cassiopea* is expected to become an ecological ‘winner’ in increasingly anthropogenically altered marine environments, exhibiting marked increases in abundance and expansion of their distribution range (Aljbour et al., 2017, 2019; Béziat & Kunzmann, 2022; Klein et al., 2019; Mammone et al., 2021; Purcell, 2012; Tilstra et al., 2022).

Disentangling the key processes in the heat stress response of *Cassiopea* would provide important insights into the ecological success of this invasive genus and improve our conceptual understanding of the mechanisms underlying thermal tolerance in cnidarian-algal symbioses in general. Specifically, we hypothesize that the nutrient status of symbiotic partners and their interaction are crucial drivers of the heat stress response of *Cassiopea* holobiont. Here we thus aimed to characterize the physiological, bioenergetic, symbiotic and cellular mechanisms involved in the gradual response and subsequent collapse of *Cassiopea* medusae holobionts during acute heat stress.

For this, we studied the symbiotic interactions and the nutritional status of the *Cassiopea* holobiont in a controlled heat stress experiment. We characterized the intrinsic response of unfed *Cassiopea andromeda* (Forskål, 1775) to prolonged exposure to elevated temperatures inducing heat stress in the holobiont (Béziat & Kunzmann, 2022; McGill & Pomory, 2008). Physiological and elemental analyses were combined with isotope labeling and correlative secondary electron microscopy (SEM) and Nanoscale Secondary Ion Mass Spectrometry (NanoSIMS) imaging to study the impact of heat stress on the metabolism of the symbiotic partners. This approach permitted the identification of similarities with well-described symbiotic responses to heat stress. It also highlighted some of the distinct characteristics potentially explaining the high heat tolerance of the *Cassiopea* holobiont.

## Materials and methods

### Animal husbandry

Forty-eight *Cassiopea* medusae with bell diameters around 2 cm (1.95 ±0.16 cm, mean ±SD) and a generally healthy appearance (intact and round bell, constant pulsation, opened oral arms) were used for this experiment. These animals were bred and reared in a culture aquarium capable of supporting the complete reproductive life cycle of *Cassiopea.* This population of animals was derived from medusae originally acquired from DeJong Marinelife in the Netherlands. Genetic identification by amplification and sequencing of fragments of the COI (mitochondrial cytochrome oxidase subunit) regions from three individuals from this culture identified the species as *Cassiopea andromeda* (data not shown). In the 200 L culture tank, the medusae were maintained in artificial seawater (ASW) prepared from sea salt (Reefs Salt, Aquaforest) at a salinity of 35 ppt and 25.3 °C ±0.2°C, illuminated with approximately 100 µmol photons m^−2^ s^−1^ from LED lights on a 12h:12 h day:night cycle. Before the experiment, the animals were fed *ad libitum* two to three times a week with freshly hatched *Artemia salina* nauplii.

### Design of the thermal stress experiment

Three days before the beginning of the experiment the selected medusae were transferred to the experimental setup for acclimatization.

The experimental setup consisted of two identical units, one for each condition, (i.e., control and heat stress), each including a main 15 L water bath, a circulation pump and a heater to maintain a homogeneous temperature (**Figure S1**). The water baths were closed by a transparent PVC cap to avoid evaporation and salinity fluctuations. Inside each water bath, six independent 500 mL transparent experimental acrylic containers were equipped with transparent lids and individual air bubbling (Loft food container, Rotho, Switzerland). Each container was filled with 400 mL of ASW, had no water exchanges with the main water bath, and hosted four medusae. Animals were kept on a 12h:12h day:night cycle with LED lights (VIPARSPECTRA V165, USA) providing approximately 110 μmol photons m^−2^ s^−1^. Each day, the salinity in the experimental containers was measured and 90 % of the ASW was replaced with fresh ASW at the same temperatures as the thermal baths. During the acclimatization and experimental period (a total of 13 days), animals were not fed to exclude potential confounding effects of treatment conditions on heterotrophic nutrient acquisition.

The temperature in both water baths was constantly recorded using a submersible temperature logger (Pendant Temperature/Light 64K Data Logger, HOBO, US) placed in a separate acrylic container containing 400 mL of ASW without medusae. For both experimental conditions, units were maintained at a temperature of approximately 27 °C (control: 27.3 ±0.5 °C, heat stress: 27.0 ±0.8 °C) during an acclimatization period of 2 days. At the beginning of the experiment (day 1), the temperature of the heat stress treatment unit was ramped up linearly at a rate of 1 °C increments every 12 h until a final temperature of about 34 °C, which was reached on day five and subsequently maintained at 34.1 ±0.5°C throughout the experiments, i.e., for seven days (**Figure 1A**). The control animals remained at an average temperature of 27.2 ±0.6 °C throughout the experiment.

**Figure 1:**
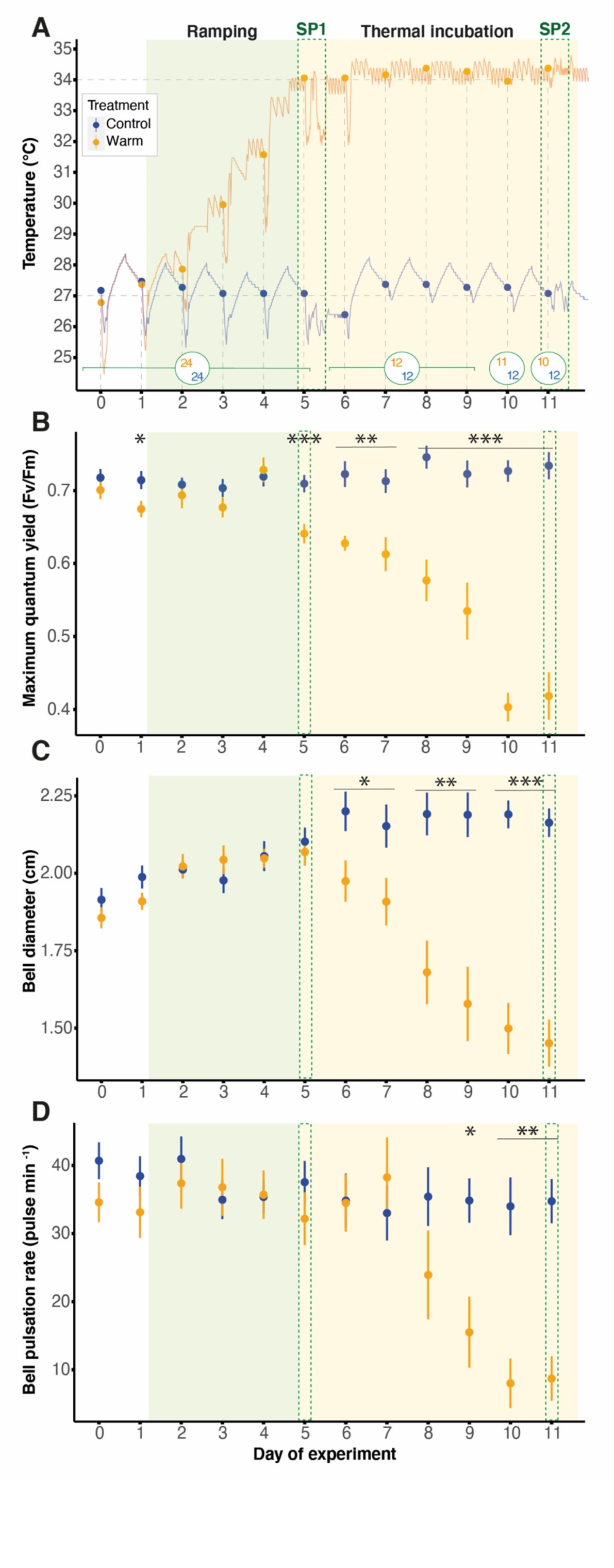
Thermal treatment and daily measurements on Cassiopea andromeda holobionts. **A** Temperature profiles of the two thermal treatments over the course of the experiment. The filled circles indicate the seawater temperature at the start of daily measurements. Numbers inside the larger open circles indicate the number of animals for each treatment at the corresponding time. **B** Maximum quantum yield (Fv/Fm) of the algal symbionts. **C** Evolution of bell diameters and **D** bell pulsation rates of the medusae over time. Filled circles and error bars indicate mean ±SE for control (blue) and heat stress (orange) treatment respectively. Asterisks indicate significant differences between treatments for respective time points (* p < 0.050, ** p < 0.010, *** p < 0.001). SP = Sampling Point.

In order to assess the impact of temperature on medusae physiology throughout the experiment, daily measurements of pulsation rate, maximum quantum yield, and bell diameter were carried out at the end of the dark period (**Figure 1**). Isotopic labeling, incubations, and sampling of water and medusae occurred on day 5 (SP1) and 11 (SP2), corresponding to the 1^st^ and the 7^th^ day at the final temperature (34 °C) of the thermal-stress treatment (**Figure 1A**).

### Daily measurements of pulsation rate, maximum quantum yield, and bell diameter

Every morning, within 1.5 h before the beginning of the light period, two sets of measurements on dark acclimated medusae were performed using low-intensity red light in order to not affect their symbiotic photophysiology. First, the night pulsation rates were assessed by counting the bell contractions of each medusae for one minute. Medusae presenting unsynchronized pulsations (spasms) were counted as not pulsating. Only medusae presenting ‘melted’ and disintegrated bell tissues were considered dead and removed from the experiment. Then, the dark-acclimated or maximum quantum yield (Fv/Fm) was measured by pulse-amplitude modulation (PAM) fluorometry using a MINI-PAM-II (blue version; Walz GmbH, Germany) with a 5.5 mm fiber optic targeting the center of the medusae.

Each day, at the beginning of the light cycle, medusae were placed on a scaled board in their experimental containers and photographed with a fully expanded bell to record their bell diameter. For this, each experimental container was removed from the water baths for less than two minutes to avoid excessive cooling of the contained water, followed by a water exchange.

### Physiological parameters

On each of the two sampling days, six medusae per condition were sampled for physiological measurements, except for the heat-stressed condition at SP2 (only four medusae were sampled due to early mortality).

To assess the ammonium (NH_4+_) uptake and release by the medusae, they were transferred individually into glass beakers containing 40 mL of freshly prepared ASW prewarmed to the respective treatment temperatures and placed inside the corresponding water baths. Additionally, one beaker containing only seawater (no medusae) was placed in each water bath to assess and correct for the potential evolution in NH_4+_ concentration in seawater unrelated to the medusa’s presence. After 6 hours of incubation in the light, the ASW from each beaker was collected and replaced with temperature-matched seawater. The collected water samples were filtered through 0.22 µm PES filters (pre-rinsed with Milli-Q water and an aliquot of sample water, which was discarded), and NH_4+_ concentrations were immediately measured using a Smartchem450 wet chemistry analyzer (AMS Alliance, Italy).

After the water sampling, the six medusae were collected to assess their wet weight, host protein content, Symbiodiniaceae density, and chlorophyll *a* content. Excess water was removed, and medusae were placed in a pre-weighted 5 mL round-bottom culture tube and weighed using a precision balance. Cold 2X PBS was added by taking into account the wet weight of the medusa and aiming for a final volume of 3 mL (Volume of PBS added = 3 mL – medusae volume, estimated from the medusae weight assuming a density of 1 g mL^-1^). The medusae were then homogenized on ice in 2X PBS using a Polytron Immersion Dispenser (Kinematica, Malters, Switzerland) for at least 30 s until the tissue slurry was visibly homogeneous. To separate the host fraction from the Symbiodiniaceae, two equal aliquots of 1.3 mL of the tissue homogenate were transferred into Eppendorf tubes and centrifuged at 3000 × g and 4 °C for 3 mins. The supernatants containing the host fraction were transferred into a 2 mL cryogenic tube, snap-frozen in liquid nitrogen, and stored at -80°C until further processing of host protein content measurements. The pellets containing the Symbiodiniaceae were resuspended in 1 mL of 2X PBS, transferred into a 2 mL cryogenic tube, snap frozen, and stored at -80 °C for subsequent Symbiodiniaceae and chlorophyll *a* quantification.

The host protein content of medusae was assessed in three technical replicates, using a Pierce Rapid Gold BCA Protein Assay Kit (ThermoFisher Scientific, Germany), following the manufacturer’s instructions for the standard microplate protocol. The host fraction of the tissue homogenate was gently thawed on ice. Technical triplicates of each sample were transferred into a flat-bottom 96-well plate. Protein content was assessed in a BioTek Synergy H1 high sensitivity plate reader (BioTek Instruments, USA) by measuring the absorbance at 480 nm and calibrated against absorbances of a serial dilution of Bovine Serum Albumin (BSA) protein standard (range of standard curve from 0 to 2 µg mL^-1^), which were assayed alongside the medusa samples.

In order to assess the density of Symbiodiniaceae, the algal symbiont pellet was gently thawed on ice, and resuspended by vigorous vortexing, and four technical aliquots of 10 µl were analyzed using a CellDrop^TM^ automated Cell Counter (DeNovix, USA). Automated cell count measurements were based on chlorophyll autofluorescence in the red channel and cell size. Finally, averaged Symbiodiniaceae counts for each medusa were standardized by host protein as a proxy of host biomass.

The chlorophyll *a* content was quantified from an aliquot of 700 µl of the Symbiodiniaceae fraction (the same aliquot from which Symbiodiniaceae cell counts were carried out). Symbiodiniaceae cells were first rinsed by two centrifugation steps (3000 × g at 4 °C for 3 and 5 mins) with a resuspension of the pellet in 1 mL of 2X PBS in between. The pellets were then resuspended in 500 µl of ice-cold ethanol (95 %) and incubated in the dark at 4 °C under constant rotation overnight (14 h) for chlorophyll extraction. Finally, 200 µl of each sample in duplicate was transferred into a flat bottom 96 well plate, along with duplicates of the solvent for the blank standardization. The absorbance of each sample was measured at 630, 664, and 750 nm with a BioTek Synergy H1 high sensitivity plate reader (BioTek Instruments, USA), and the chlorophyll *a* content was calculated as follows (with turbidity correction, Jeffrey and Humphrey,1975):

Chlorophyll *a* (µg mL^-1^) = 11.43*(OD(664)-OD(750))-0.64*(OD(630)-OD(750))

Chlorophyll *a* content was then standardized to the host protein content of each individual medusa.

### Elemental composition

In order to measure the impact of temperature on the total organic carbon and nitrogen content of the medusa, three individuals per treatment and sampling time point were sampled. After seawater was carefully removed, the medusae were weighted in a pre-weighed 5 mL round-bottom culture tube, and a volume of 2X PBS was added to a final volume of 3 mL, as described above. The tissue was then homogenized, aliquoted, and the host and dinoflagellate fractions were separated and snap frozen with an additional rinsing step for the Symbiodiniaceae prior to snap freezing in liquid nitrogen (3000 × g at 4 °C for 3 mins, and resuspension of the pellet in 1mL of 2X PBS). Frozen host and Symbiodiniaceae fractions, were freeze-dried for two days. Approximately 2.5 ±0.4 and 10.5 ±1.5 mg of the respective dried sample fractions were then weighed in duplicate for each sample and encapsulated in aluminum for carbon and nitrogen analysis respectively. The elemental analysis/isotope ratio mass spectrometry (EA/IRMS) was performed using a Carlo Erba 1108 (Fisons Instruments, Italy) elemental analyzer connected via a ConFlo III split interface to a Delta V Plus isotope ratio mass spectrometer (Thermo Fisher Scientific, Germany) operated under continuous helium flow (Spangenberg & Zufferey, 2018). The carbon and nitrogen contents (in wt. %) were determined from the EA/IRMS peak areas and converted into the atomic abundance to calculate atomic C:N ratios for each sample.

### Stable isotope labeling experiment for nutrient tracking and histological analyses

In order to investigate the impact of the temperature on the uptake and exchange of nutrients between the medusae and the Symbiodiniaceae, three medusae were incubated in isotopically labeled ASW for each condition and sampling point.

One day before labeling and sampling, ASW was freshly prepared with Milli-Q water, depleted of dissolved organic carbon by acidification with HCl (4 M) to a pH < 3, and maintained under constant bubbling with air for at least 4 h. This ASW was then supplemented with ^13^C-bicarbonate (Sigma 372382) to a final concentration of 3 mM, and the pH of the solution increased again to 8.1 with 1 M NaOH solution. Finally, ^15^N-ammonium chloride (Sigma 299251) was added to a final concentration of 3 µM. After thorough mixing, the labeled ASW was split into two aliquots and thermally equilibrated overnight at the temperatures for the control and heat stress treatments, respectively.

Incubation with isotopically labeled bicarbonate and ammonium was initiated at the beginning of the light period by placing three medusae into glass beakers filled with 40 mL of thermally equilibrated isotopically-labeled ASW medium for 6 h. After three hours into this incubation, 30 mL of isotopically labeled ASW medium was replaced to ensure constant concentrations of ^13^C-bicarbonate and ^15^N-ammonium, respectively. At the end of the incubation, the medusae were sampled and cut into quarters using clean razor blades. Two of these quarters were fixed immediately for correlative SEM and NanoSIMS imaging (4 % paraformaldehyde, 2.5 % glutaraldehyde in 0.1 M Sorensen’s buffer, 9 % sucrose), and one quarter was fixed for paraffin embedding and light microscopy (4 % paraformaldehyde in 0.1 M Sorensen’s phosphate buffer pH 7.9, 9 % sucrose). Fixed samples were maintained at 4 °C until further processing. To have an unlabeled control sample, one medusa within the same size range was sampled directly from the culture tank, dissected, preserved, and processed in the same manner.

### Nutrient assimilation and cellular ultrastructure by correlative SEM and NanoSIMS imaging

The bell tissue samples fixed for resin embedding (for correlative SEM and NanoSIMS imaging) were processed after at least one night of fixation at 4 °C. First, the samples were rinsed with Sorensen’s buffer (0.1 M) to remove the fixative and dissected to obtain a small piece of tissue (around 4 mm^3^) from the center of the bell. To preserve the lipid fraction of the samples, the small tissue pieces were post-fixed for 1 hour with osmium tetroxide (OsO_4_ 1 %, 1.5 % potassium hexacyanoferrate II in 0.1 M Sorensen phosphate buffer) under constant agitation and rinsed twice in Milli-Q water for 20 minutes. Using a tissue processor, the samples were then subjected to a serial dehydration in ethanol (30 %, 70 %, and 100 % ethanol in Milli-Q water), to facilitate a progressive Spurr resin infiltration (30 %, 70 % and 100 % Spurr resin in absolute ethanol). Once infiltrated, the samples were placed into molds filled with 100 % Spurr resin and cured at 60 °C for 48 hours. Thin sections (200 nm) were cut from the resin blocks using an Ultracut S microtome (Leica Microsystems) and a diamond knife (DiATOME) and collected on clean silicon wafers.

In order to add contrast and to visualize the subcellular structures present in the tissues, sample sections were post-stained with 1 % uranyl acetate and Reynolds Lead Citrate before imaging by scanning electron microscopy (SEM, GeminiSEM 500, Zeiss; 3 kV, aperture size of 30 µm, and a working distance of 2.9 to 2.3 mm) with an energy selective backscatter detector (EsB, grid of 130 V; Zeiss).

To image the distribution of isotopic enrichments within subcellular structures visualized in SEM images, the same sections were sputter coated with a 12 nm gold layer (using a Leica EM SCD050 gold coater) and transferred to a Nano Secondary Ion Mass Spectrometry (NanoSIMS) 50 L for analysis (Hoppe et al., 2013). In the NanoSIMS, the pre-sputtered samples were bombarded with a Cs^+^ primary ion beam at 16keV with a current of around 2 pA, focused to a spot size of ca. 150 nm. This beam was rastered over an area of 45 × 45 µm with a dwelling time of 5000 µs per pixel. The secondary ions ^12^C^12^C^-^,^12^C^13^C^-^,^12^C^14^N^-^,^12^C^15^N^-^ were counted individually in electron multiplier detectors at a mass resolution power of around 9000 (Cameca definition), which resolves potential interferences in the mass-spectrum. From the resulting isotope maps (45×45 µm^2^, 256×256 pixels, superposition of 6 drift-corrected images), regions of interest (ROIs) were drawn around the different tissue compartments, i.e., Symbiodiniaceae, amoebocytes (excluding the Symbiodiniaceae), and the epidermis. For each ROI, the isotopic ratio enrichments established through the ratios ^12^C^13^C^-^/^12^C_2-_ and ^15^N^12^C^−^/^14^N^12^C^−^ were quantified against a control sample with natural isotopic compositions prepared and analyzed in an identical manner, using the NanoSIMS software L’Image (v.10-15-2021, developed by Dr. Larry Nittler). Isotope enrichments are reported in the delta (δ) notation as followed:

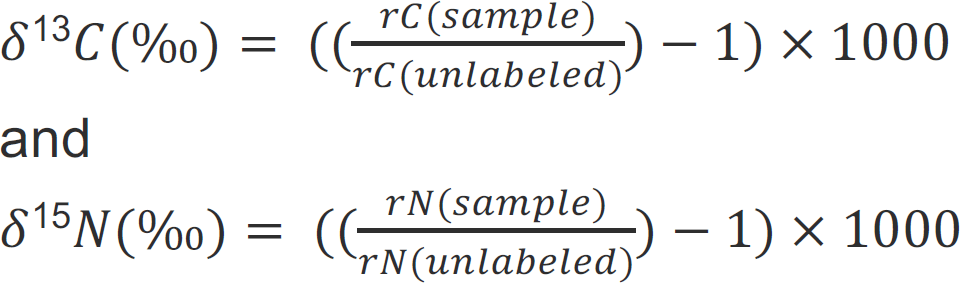

where rC_(sample)_ and rC_(unlabeled)_ are the count ratios of ^12^C^13^C^-^/^12^C_2-_ in the sample and in an unlabeled control, respectively. rN_(sample)_ and rN_(unlabeled)_ are the count ratios of _15_N^12^C^−^/^14^N^12^C^−^ in the sample and in an unlabeled control, respectively.

Compartments were only considered to be significantly enriched if the average of the delta value of the measured ROIs were more than two standard deviations above the average delta values measured in similar compartments in an unlabeled sample. The number of images and ROIs per compartment and sample is reported in the supplementary information (**Table S1**).

In order to estimate the symbiont contribution to host metabolism, the mean of ratios of ^13^C enrichments in host amoebocytes and their algal symbionts was calculated: each ^13^C enrichment value of an amoebocyte was divided by the mean of the ^13^C enrichments of the symbionts hosted within this amoebocyte. Only viable amoebocytes and symbiont cells that were significantly enriched were included in this calculation.

### Histological characterization of the tissue structure and cell density in the medusae bell tissue by light microscopy

The bell tissue fixed for paraffin embedding underwent serial dehydration in ethanol, then xylene before infiltration and embedding in paraffin. Sections 4 µm thick of paraffin-embedded samples were cut and placed on a glass slide, dewaxed, and stained with hematoxylin and eosin (H&E) before being imaged by transmitted light microscopy with a Keyence VHX-7000 digital microscope. The number of host nuclei (stained in blue by the H&E staining) and the number of dinoflagellate cells in the mesoglea were assessed and standardized by the mesoglea surface area using ImageJ2/Fiji plugin Cell counter and Measure (v2.9.0, https://github.com/fiji/Cell_Counter/blob/Cell_Counter-3.0.0/src/main/java/sc/fiji/cellCounter/CellCounter.java) in order to obtain cell density. Two different areas of the mesoglea were assessed on the images: the oral mesoglea, located between the oral epidermis and the gastrodermis and hosting most of the dinoflagellates, and the inner mesoglea, located below the gastrodermal tissue layers and hosting few dinoflagellates. This quantification was performed on three images per sample, and on three samples per condition (all with the same settings and same magnification).

### Statistical Analysis

All analyses were performed in R (version 4.2.0, (Venables & Smith, n.d.). All response parameters were measured daily among treatments and were analyzed for each experimental day using ANOVA (for normally distributed data, considered true when *p* < 0.05 for Shapiro-Wilk test) or Kruskal-Wallis (when the normal distribution assumption of data was violated), followed by multiple comparison corrections of p-values based on false discovery rate (FDR).

For the physiological measurements, the influence of the treatment and sampling point on the data were analyzed using a two-factorial ANOVA (normal distribution of the data as confirmed by Shapiro-Wilk test, *p* > 0.05) followed by Tukey’s honestly significant differences (HSD) post hoc comparison. In the cases where ANOVA results indicated a significant interaction between treatments and sampling points but Tukey’s HSD did not allow resolving treatment effects for individual sampling points, one-factorial ANOVA was used for individual sampling points (indicated in the figures with asterisks in brackets). The ammonium uptake data were square-root transformed following the addition of a constant to avoid negative values before analysis to assure the homogeneity of the variances required for the ANOVA. The isotopic enrichment across treatments and sampling points was analyzed with a linear mixed model (LMM) introducing the three biological replicates as a random variable. This was followed by a Tukey’s HSD post hoc comparison. In the cases where LMM results indicated a significant interaction between treatments and sampling points but Tukey’s HSD did not resolve treatment effects for individual timepoints, one-factorial ANOVA or Kruskal-Wallis was used for individual timepoints (indicated in the figures with asterisks in brackets).

Finally, differences in cell density in the mesoglea among treatments for sampling point 2 were analyzed with a linear mixed model (LMM), using the three biological replicates as a random variable. All data in the text are presented as mean ±SD unless stated otherwise.

## Results

### Response of Cassiopea andromeda to elevated seawater temperature

During the ramping period of the experiment (**Figure 1A**), the increase in seawater temperature had no detectable effect on the physiology of the medusae (**Figure 1B-D**). However, once the maximum temperature (34 ºC) was reached on day 5, the *Cassiopea* holobionts started to show the first response to heat stress compared to the control specimens. While the maximum quantum yield (Fv/Fm) of algal symbionts remained stable over time under control conditions, their heat-stressed counterparts showed a significant decline starting from day 5 of the experiment (ANOVA, F = 14.61, FDR-adjusted *p* < 0.001). This decline became increasingly pronounced over time, leading to a 43 % reduction in heat-stressed (Fv/Fm = 0.42 ± 0.10) compared to control (Fv/Fm = 0.73 ± 0.06) holobionts on the last day of the experiment (ANOVA, *F* = 77.19, FDR-adjusted *p* < 0.001, **Figure 1B**).

In addition to the photophysiological response of the algal symbionts, host metrics were visibly affected by the heat stress treatment. While the bell diameters of medusae tended to increase similarly in both treatments during the ramping period, the bell diameters of heat-stressed medusae exhibited a significant decrease beginning on day 6 (ANOVA, *F* = 5.98, FDR-adjusted *p* = 0.046; **Figure 1C**), resulting in an average 33 % decline compared to control medusae by the end of the experiment. Likewise, the pulsation rate in heat-stressed medusae decreased significantly from day 9 onwards (ANOVA, *F* = 10.28, FDR adjusted *p* = 0.016) with a 75 % decline by the end of the experiment (**Figure 1D**). This drop in pulsation rate in heat-stressed medusae was also associated with spasms and/or total immobility in some individuals.

Ultimately, the first cases of mortality were observed in heat-stressed medusae towards the end of the experiment, beginning on day 10 (**Figure 1A**).

### Heat stress alters the physiology and the nutritional status of Cassiopea andromeda

The *Cassiopea* host exhibited a clear physiological response to heat stress already a few days after reaching 34 °C (i.e., around days 6 and 7 of the experiment; **Figure 1**) and this trend continued until the end of the experiment, resulting in a number of pronounced effects between the two sampling points SP1 (day 5) and SP2 (day 11) (**Figure 2**).

**Figure 2:**
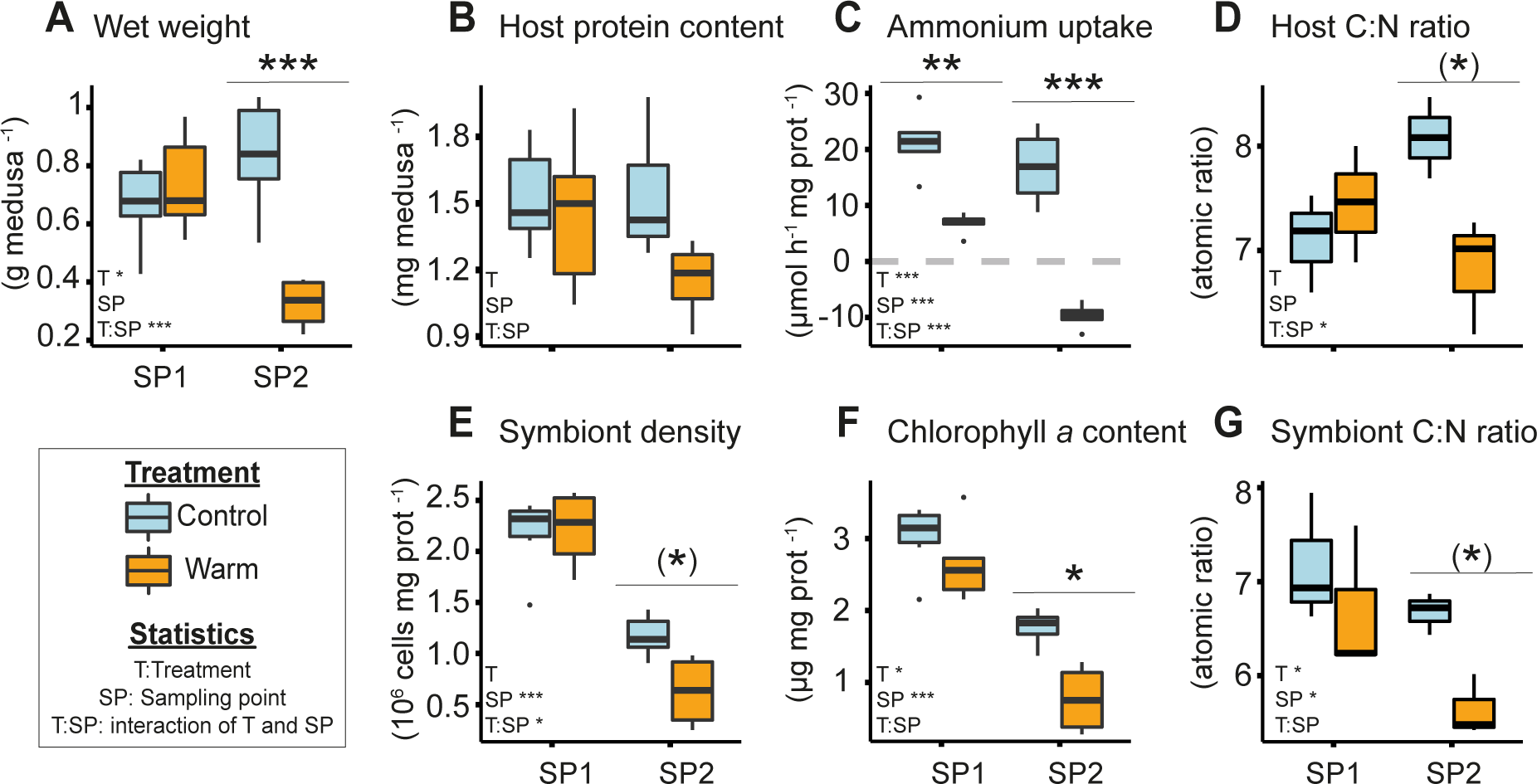
Physiology and nutritional status of Cassiopea holobionts exposed to heat stress. **A** Medusae wet weight, **B** host protein content, **C** holobiont net ammonium uptake, **D** host atomic C:N ratio, **E** symbiont density per host protein, **F** chlorophyll a content per host protein, and **G** symbiont atomic C:N ratio. Sampling points correspond to day 5 (SP1) and 11 (SP2) of the experiment. Asterisks indicate significant differences between treatments (* p < 0.050, ** p < 0.010, *** p < 0.001). Asterisks in brackets show significant differences when only SP2 is considered. Number of biological replicates per condition: **A,B,C,E,F** n = 6 except for SP2 under heat stress where n = 4; **D,G** n = 3.

The wet weight of the medusae was unchanged at SP1 (Tukey’s HSD, *p* = 0.891), but showed a significant 61 % decrease at SP2 under thermal stress (Tukey’s HSD, *p* < 0.001; **Figure 2A**). The host protein content per medusa exhibited no significant effect of heat stress at SP1, but a 25 % reduction (albeit not significant) at SP2 (ANOVA, *F* = 2.87, *p* = 0.107; **Figure 2B**). Net ammonium uptake rates were down by a significant 51 % decrease under heat stress already at SP1 (Tukey’s HSD, *p* = 0.009). They even showed a negative rate (i.e., a net release of ammonium into the surrounding seawater) at the end of the experiment (Tukey’s HSD, *p* < 0.001, **Figure 2C**). The C:N ratios of heat-stressed host medusae remained stable at SP1 (Tukey’s HSD, *p* = 0.828), but exhibited a significant 16 % decrease at SP2 (Tukey’s HSD, *p* = 0.059). Interestingly, the average host C:N ratio of the control animals showed a non-significant 14 % increase over the course of the experiment (Tukey’s HSD, *p* = 0.154, **Figure 2D**).

Despite these changes in host physiology, medusae showed no visual signs of bleaching, and no loss of pigmentation was observed during the experiment (**Figure S3**). Nonetheless, the physiology and elemental compositions of the symbionts were clearly affected by heat stress at both sampling points (**Figure 2E-G**). The density of symbiont cells relative to host protein content in heat-stressed animals decreased significantly by 46 % at SP2 (ANOVA considering only SP2, *F* = 9.54 *p* = 0.015; **Figure 2E**). The chlorophyll *a* content of symbionts (per mg of host protein) became significantly reduced during heat stress (ANOVA, *F* = 7.29, *p* = 0.015) with a 55 % decline compared to control conditions at SP2 (Tukey’s HSD, *p* = 0.011, **Figure 2F**) and the symbiont C:N ratio was significantly reduced during heat stress, down by 16 % at SP2 (ANOVA, *F* = 5.55, *p* = 0.046) compared to control conditions (ANOVA considering only SP2, *F* = 20.66, *p* = 0.011, **Figure 2G**). However, when normalized to wet body weight, neither symbiont densities nor chlorophyll *a* content was significantly affected by heat stress (symbiont densities per wet weight: LMM, *F* = 0.02; *p* = 0.894; chlorophyll *a* per wet weight: LMM, *F* = 0.45, *p* = 0.513; **Figure S3**).

### Heat stress reduces nutrient assimilation in the symbiosis

Overall, NanoSIMS analyses revealed that heat stress had pronounced effects on ^13^C-bicarbonate and ^15^N-ammonium assimilation by the symbiotic partners (**Figure 3**).

**Figure 3:**
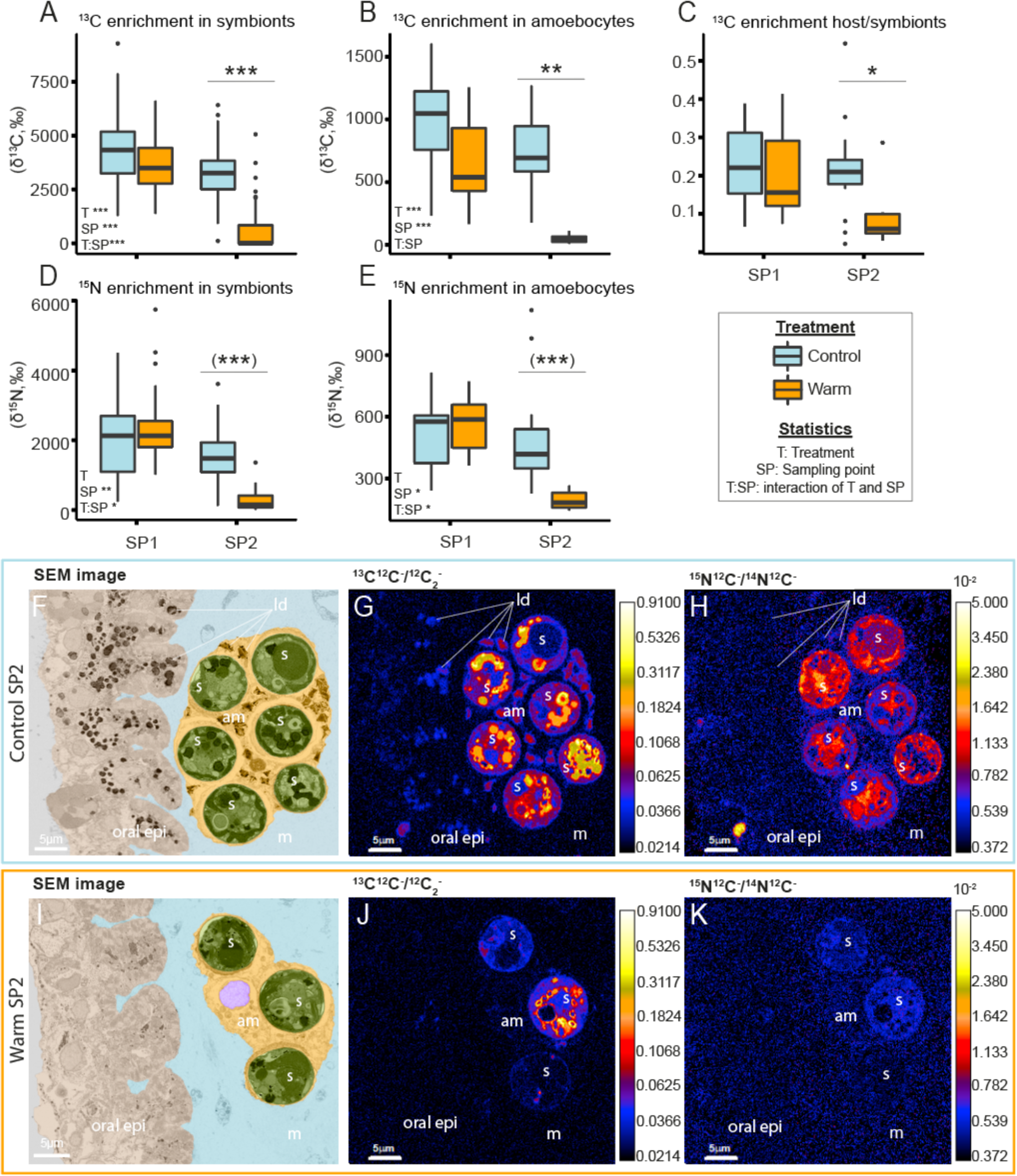
Temperature effects on enrichment and (sub)cellular localization of isotopically labeled carbon and nitrogen in heat-stressed medusae. **A** ^13^C enrichment from ^13^C-bicarbonate (H^13^CO_3_^-^) assimilated into algal symbiont cells and B translocated to their host amoebocyte cells. C Ratio of ^13^C enrichment between host amoebocytes and their algal symbionts. D ^15^N-enrichment induced by the assimilation of ^15^N-ammonium (NH_4_+) into algal symbiont cells and E into host amoebocytes. Asterisks indicate significant differences between treatments (* p < 0.050, ** p < 0.010, *** p < 0.001). Asterisks in brackets show significant differences when only SP2 is considered. F-K Correlative SEM (F,I) and NanoSIMS (G,H,J,K) isotope ratio images of medusa sections under control conditions (F-H) and heat stress (I-K) at SP2. The SEM images are artificially colored, with oral epidermis (oral epi) in beige, mesoglea (m) in blue, amoebocyte (am) in yellow, amoebocytes nuclei in purple, symbionts (s) in green and lipid droplets (ld). The color scales of the NanoSIMS images are logarithmic.

Compared with the control condition, heat stress caused a significant reduction in ^13^C enrichment in the algal symbionts (LMM, *X^2^* = 64.54, *p* < 0.001, **Figure 3A**), in the host amoebocyte cells containing them (LMM, *X^2^* = 24.86, *p* < 0.001, **Figure 3B**), and in the host epidermis (LMM, *X^2^* = 7.30, *p* = 0.006, **Figure S4A**). In more detail, at SP1, the ^13^C enrichment of amoebocytes was insignificantly lower in heat-stressed animals (Tukey’s HSD, *p* = 0.189), while ^13^C enrichment of the algal symbionts remained relatively stable (Tukey’s HSD, *p* = 0.115). At SP2, ^13^C enrichment of both amoebocytes and symbionts under heat stress declined by 94 % and 82 %, respectively (Host: Tukey’s HSD, p = 0.006; symbiont: Tukey’s HSD, *p* < 0.001; **Figure 3A,B**). Furthermore, the average of the ratios of ^13^C enrichments in amoebocytes and their algal symbionts (excluding algae-amoebocyte pairs without significant enrichment), was reduced by 60 % (ANOVA, *F* = 6.57, *p* = 0.013) compared to control conditions (Tukey’s HSD, *p* = 0.012, **Figure 3C**).

These quantitative changes in ^13^C enrichment were accompanied by marked differences in the subcellular spatial distribution within the tissues of heat-stressed medusae at SP2 (**Figure 3F-K**). In the host tissue of control animals, strong ^13^C enrichments were concentrated in hot spots corresponding to lipid droplets (dark structures stained by osmium in SEM images, **Figure 3F,G**). During heat stress, however, these lipid droplets appeared less abundant in the epidermis and almost absent in the amoebocytes, and thereby likely contributing to a lower ^13^C enrichment in the host (**Figure 3I,J**).

In the symbionts, ^13^C enrichments were primarily located in the pyrenoid and lipid droplets under control conditions (**Figure 3H**). Under heat stress, however, ^13^C enrichment hotspots were overall less pronounced in symbiont cells (**Figure 3K**) with some cells exhibiting almost no detectable ^13^C enrichment.

While heat stress also affected ^15^N-ammonium assimilation by the symbiotic partners, the extent was less pronounced. Heat stress caused no overall significant effect on _15_N enrichment due to ammonium assimilation in the host epidermis (LMM, *X^2^* = 0.37*, p* = 0.541, **Figure S4B**), the amoebocyte cells (LMM, *X^2^ =* 2.07, *p* = 0.150, **Figure 3E**), and the algal symbionts they contained (LMM, *X^2^* = 2.57, *p* = 0.109, **Figure 3D**). Furthermore, the interaction of the sampling time and heat stress factors had no significant effect on the reduction of ^15^N enrichment of the host epidermis (LMM, *X^2^* = 0.41, *p* = 0.521), but had an impact on the amoebocytes (LMM, *X^2^* = 5.77, *p* = 0.016) and algal symbionts (LMM, *X^2^* = 5.12, *p* = 0.024). When analyzed per time point, the _15_N enrichment in the amoebocytes and symbionts of heat-stressed medusae remained stable compared to the control treatment at SP1 (Amoebocyte: Kruskal Wallis considering only SP1, *X^2^* = 1.85, *p* = 0.172; symbiont: Kruskal Wallis considering only SP1, *X^2^* = 2.43, *p* = 0.119), but exhibited significant decreases (60 % and 84 %, respectively) at SP2 (Amoebocyte: Kruskal Wallis at SP2, *X^2^* = 23.86, *p* < 0.001; symbiont: Kruskal Wallis at SP2, *X^2^* = 87.09, *p* < 0.001). This was accompanied by differences in the subcellular spatial distribution of ^15^N-ammonium enrichment between temperature treatments at SP2 (**Figure 3I-K**). Specifically, ^15^N-ammonium enrichment was largely homogenous in the host tissues, with strong enrichment in the symbionts under control conditions (**Figure 3F,H**). During heat stress, however, ^15^N enrichment was less pronounced in the host tissue and substantially reduced in the symbionts, with some cells showing no discernible enrichment (**Figure 3I,K**).

### Impact of temperature on tissue and cells ultrastructure

Light and electron microscopy of the bell tissues revealed an increase in the density of host cells in the mesoglea in heat-stressed medusae at SP2. This was most pronounced in the oral mesoglea and was accompanied by substantial *in hospite* degradation of algae cells (**Figure 4, S5**).

**Figure 4:**
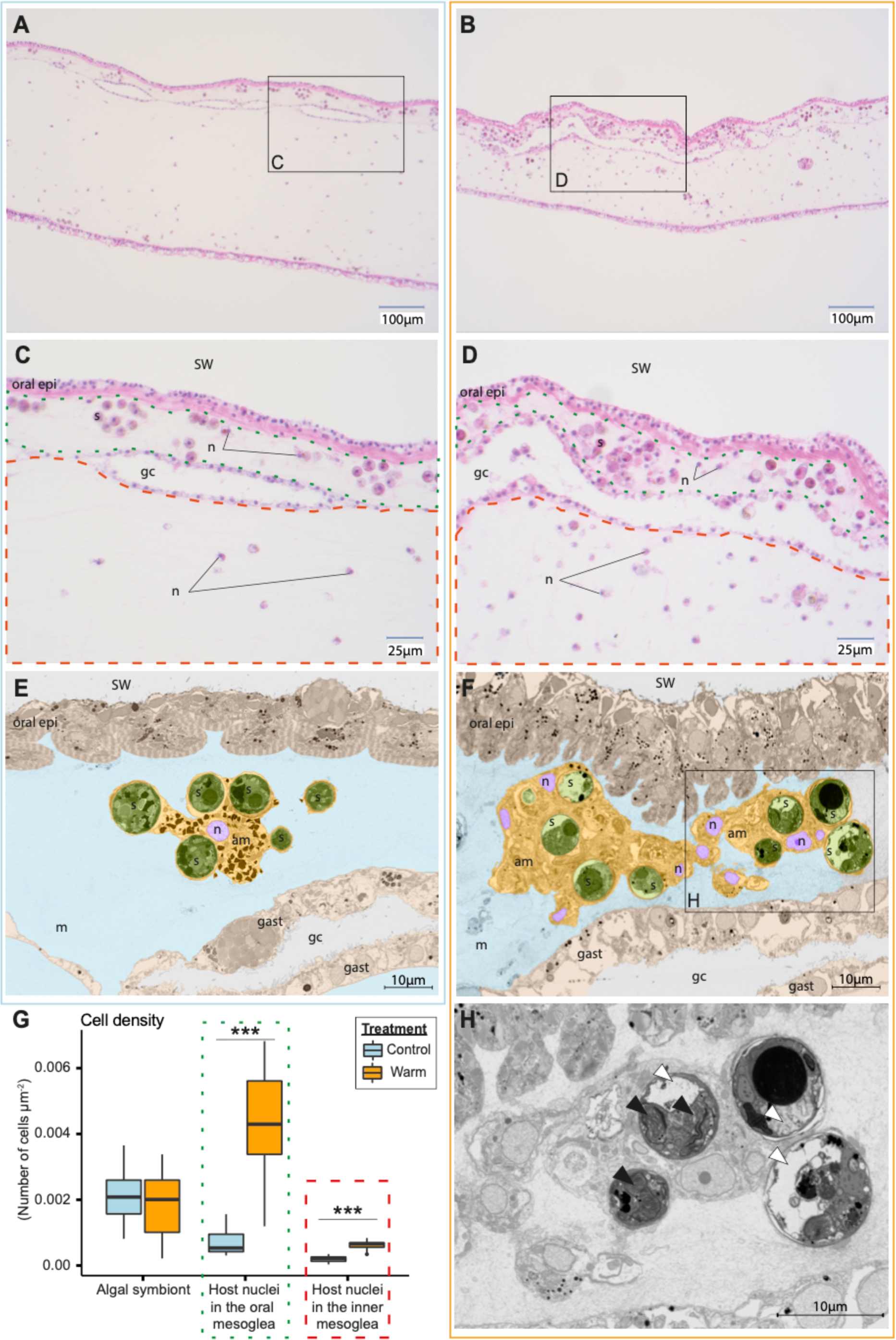
Temperature effects on the ultrastructure and cell density of the bell tissue of Cassiopea andromeda medusae. A-D Light microscopy images and E,F,H SEM images of the medusae in control (A,C,E, i.e., left column) and heat stress conditions (B,D,F,H, i.e., right column). E and F are artificially colored for better visualization with oral epidermis and gastrodermis in beige, mesoglea in blue, amoebocyte(s) in yellow, amoebocyte nuclei in purple, and symbionts in green. G Impact of heat stress on the density of symbionts and host nuclei in the oral (delineated by green dotted line) and inner mesoglea (delineated by red dashed line) illustrated in C and D. Asterisks indicate significant differences between treatments (* p < 0.050, ** p < 0.010, *** p < 0.001). H SEM image exhibiting the intracellular degradation (white arrowhead) and disorganized thylakoids (black arrowhead) in heat-stressed symbionts. SW: seawater, oral epi: oral epidermis, m: mesoglea, am: amoebocyte(s), n: amoebocytes nuclei, s: symbionts, gast: gastrodermis).

Light microscopy demonstrated around six- and threefold increases in the densities of host nuclei for the oral (LMM, *X^2^* = 12.16, *p* < 0.001) and inner mesoglea, respectively (LMM, *X^2^* = 35.44, *p* < 0.001, nuclei colored in dark blue by the H&E staining in **Figure 4A-D**, quantified in **Figure 4G**). In contrast, the density of algal symbionts in the oral mesoglea remained unchanged by the heat treatment (LMM, *X^2^* = 0.09, *p* = 0.759). SEM imaging corroborated the observation made in light microscopy, with a high number of host nuclei in the oral mesoglea of heat-stressed medusae (**Figure 4F**). These host nuclei appeared to be mostly in clustered groups of host cells, some of which were hosting algal symbionts. In contrast, such host cell clustering was not visible in the tissues of control medusae (**Figure 4E**). The cellular membranes of each cell in these clusters were difficult to visualize, not permitting any interpretation of their health state, apart from the absence of visible nuclei condensation. Importantly, under heat stress, the symbiotic algae present in these amoebocytes displayed clear signs of stress with different degrees of deterioration. Specifically, these symptoms included disorganization of cellular contents and thylakoids and pronounced internal degradation of cell contents (**Figure 4H**).

## Discussion

Climate change has resulted in an unprecedented decline of tropical coral reef ecosystems at a global scale due to the demise of their main ecosystem engineers, reef-building corals. In corals, this decline is partly linked to the breakdown of symbiotic nutrient cycling, leading to host starvation that ultimately results in coral bleaching and mortality (Cunning et al., 2017a; Morris et al., 2019; Rädecker et al., 2021). In contrast, the upside-down jellyfish *Cassiopea* is projected to thrive under the current and future conditions despite hosting similar photosynthetic symbionts (Purcell, 2012). Here, we were able to show that severe heat stress enhanced host catabolism and reduced the contribution of the algal symbionts to the metabolism of *Cassiopea*, creating a state of host starvation in unfed animals. While this stress response resembles that observed in corals, our data suggest that some mechanistic aspects of the breakdown of the symbiosis reflect the unique properties of the *Cassiopea* holobiont. Specifically, a high occurrence of *in hospite* degradation of amoebocyte-hosted algal symbionts coupled with shrinkage (i.e., a reduction in animal body size and wet weight) resulted in a delayed and ‘invisible’ bleaching response.

### Heat-induced host catabolism and carbon starvation

In this study, severe heat stress caused host energy limitation and the metabolic switch towards catabolism, resulting in a significant depletion of carbon reserves and a visible shrinkage of *Cassiopea* medusae.

We observed a pronounced reduction of C:N ratios in heat-stressed *Cassiopea* (**Figure 2D**) coupled with a reduction of lipid droplets within the amoebocytes (**Figures 3G,J and S5**) and a less pronounced decrease in protein content (**Figure 2B**). Together, these responses point to an enhanced consumption and depletion of carbon reserves in the host metabolism. The metabolic activity of ectothermic animals tends to increase with temperature (Gillooly et al., 2001). Acute heat stress thus stimulates the respiratory energy demand of Cnidaria, including scyphozoan medusae (Aljbour et al., 2017; W. K. Fitt & Costley, 1998; Nevarez-Lopez et al., 2020). Our results suggest that *Cassiopea* consumed its energy reserves in response to such a heat-induced increase in energy demand. This is consistent with a previous study documenting the consumption of carbon reserves, more specifically of glucose and glycogen reserves by the aposymbiotic scyphozoan medusa *Stomolophus meleagris* during heat stress (Nevarez-Lopez et al., 2020).

In addition to this enhanced consumption of sugar and lipid reserves, the progressive metabolic switch from anabolism to catabolism under heat stress was also directly reflected in the nitrogen metabolism of the holobiont. The measured decrease in net ammonium assimilation at SP1 and the net release of ammonium at SP2 (**Figure 2C**) coupled with a (non-significant) decrease in host protein content (**Figure 2B**) indicate that the host gradually shifted towards a catabolic degradation of protein reserves during heat stress. In an anabolic state, cnidarian hosts utilize carbon backbones for amino acid synthesis by assimilating ammonium. In a catabolic state, proteins and amino acids are used as a carbon source for energy metabolism resulting in the production of excess ammonium by the host (Haunerland, 2003). Our findings suggest that the increased energy requirements during heat stress may have resulted in the gradual consumption of glycogen and lipid reserves and, eventually, a breakdown of proteins in the host.

Importantly, this catabolic consumption of host biomass was directly reflected in the phenotype and behavior of the animals during heat stress. Specifically, the observed reductions in bell diameter (**Figure 1C**) and wet weight (**Figure 2A**) align with previous reports documenting reductions in medusae size and weight in response to heat stress (Béziat & Kunzmann, 2022; Klein et al., 2019; McGill & Pomory, 2008; Nevarez-Lopez et al., 2020). In light of host carbon starvation during heat stress, we propose that this shrinkage is predominantly driven by the catabolic degradation of the mucopolysaccharide and collagen matrix of the mesoglea. The loss of these structural sugars and proteins, together with their ability to retain water, likely caused water loss and consequently body shrinkage in heat-stressed medusae (Chapman, 1953; Gardner & Zubkoff, 1978; Pedersen & Vilgis, 2019). This hypothesis is corroborated by reports of similar rapid change in the water content of *Cassiopea* exposed to thermal stresses (Aljbour et al., 2017) and similar shrinkage of *Cassiopea* and *Aurelia aurita* ephyrae during starvation without heat stress (Fu et al., 2014; Muffett et al., 2022). Furthermore, the observation of reduced pulsation (**Figure 1D**) and the onset of host mortality (**Figure 1A**) implies that our unfed *Cassiopea* animals were unable to compensate for the metabolic constraints of severe heat stress over extended periods of time, here 7 days.

### Disturbance of symbiotic metabolic exchange during heat stress

In contrast to heat-stressed *Cassiopea*, the animals in the control condition (i.e., 27 °C) showed no signs of carbon limitation. Even in the absence of heterotrophic feeding, these animals continued to grow over the course of the experiment (**Figures 1C, 2A, S2**). Increased C:N ratio (**Figure 2D**) and a continuous net ammonium uptake (**Figure 2C**) indicate that the release of photosynthates by the algae was sufficient to fulfill the energetic carbon requirements and to support host anabolism under control conditions. Under heat stress however, host carbon starvation and body shrinkage, together with the decline in anabolic ^13^C assimilation observed by NanoSIMS, corroborated the notion that algae-derived organic carbon was insufficient to fulfill the energetic requirements of the medusae (**Figure 3A,B**). This observed reduction in ^13^C enrichment could result from the reduced assimilation and translocation and/or enhanced catabolic consumption of metabolites during heat stress. Given that both symbiotic partners showed this trend (i.e., a reduction in ^13^C enrichment between SP1 and SP2), we suggest that heat stress reduced the relative contribution of photosynthates to the metabolism throughout the *Cassiopea* holobiont. In addition, we observed a decrease in the ratio of ^13^C enrichments in amoebocytes over that of their symbionts during heat stress (**Figure 3C**). Similar carbon retention, promoted by a decreased production and availability of photosynthates in the algal symbionts, has also been described in other symbiotic cnidarians (Baker et al., 2018; Rädecker et al., 2018).

Overall, this notion of reduced carbon availability in the symbiosis during heat stress is supported by the observed patterns of ^15^N assimilation (**Figure 3D,E**). The reduction in ^15^N assimilation by both symbiotic partners during heat stress likely reflects the combined consequences of increased catabolic production of ’unlabeled’ ammonium in the host metabolism and reduced availability of carbon backbones for ammonium assimilation (Cui et al., 2022).

Taken together, our results suggest that heat stress shifted the *Cassiopea* holobiont from a nitrogen-limited to a carbon-limited state. Reduced fixation of photosynthetic carbon and/or enhanced catabolic consumption of host carbon content (sugars, lipids, and amino acids) coincided with the enhanced retention of photosynthates by the algae. In addition, heat stress might not only deprive the host of the nutritional benefit of harboring algal symbionts, but likely also imposes additional energy requirements on the host to mitigate the harmful impacts of hosting stressed algae (e.g., the release of reactive oxygen species).

Heat stress thus effectively undermines the ecological benefits of harboring algal symbionts for the cnidarian host and might even turn the symbiosis into an additional energetic burden.

### “Invisible bleaching” by *in hospite* symbiont degradation

In contrast to the well-described bleaching response in corals, there was no evident bleaching in heat stressed *Cassiopea* in the present study (**Figure S3**). Nonetheless, a marked drop in the maximum quantum yield of algal symbionts (**Figure 1B**), a decline in the density of symbiont cells and chlorophyll *a* content normalized to host protein (**Figure 2E,F**), and some host mortality were observed under heat stress (**Figure 1A**). Hence, the absence of visible bleaching does not imply the absence of a symbiotic breakdown in the *Cassiopea* holobiont. Rather, the loss of algal symbionts was likely compensated by the shrinkage of heat-stressed medusae due to host starvation and water loss, as indicated by stable algal symbiont densities and chlorophyll *a* when normalized to wet weight (**Figure S3**). Notably, however, bleaching events characterized by a visible decrease in pigmentation of *Cassiopea* have been described previously (Klein et al., 2019; McGill & Pomory, 2008). Hence, the absence of visible bleaching in the present study reveals that the heat stress response of *Cassiopea* may depend on the species, environmental and rearing conditions, life stage, and especially the nutritional status of the medusae. The fact that the medusae were unfed during our study may have accelerated host starvation and the associated body shrinkage, thus concealing the concomitant loss of symbionts, compared to other studies. The ‘invisible’ bleaching phenomenon observed here might partially explain the low number of bleaching events documented for *Cassiopea* in the wild (Djeghri et al., 2019; Klein et al., 2019).

In corals and most other symbiotic Cnidaria, bleaching involves a multitude of mechanisms of symbiont loss, including expulsion, *in hospite* degradation, and host cell detachment (Weis, 2008). Among these mechanisms, previous studies suggested that symbiont expulsion is the most important pathway of symbiont loss during bleaching (Bieri et al., 2016). While the presence of symbionts in the gastrodermis suggests that algal expulsion occurred, our experiment did not allow us to assess the importance of this mechanism in heat-stressed *Cassiopea*. However, our ultrastructural observations suggest that other mechanisms strongly contributed to symbiont loss in *Cassiopea*. Specifically, the high abundance of heavily damaged algal symbiont cells in the gastrodermal tissues (**Figure 4F,H, S5**) combined with the drop in algal nutrient assimilation (**Figure 3A,D**) suggests a high occurrence of *in hospite* degradation of symbionts (Franklin et al., 2004). Contrary to corals and sea anemones, algal symbionts in scyphozoan medusae reside in amoebocyte host cells within the mesoglea, without direct contact with the gastrovascular cavity of the animal (Colley & Trench, 1985; Djeghri et al., 2019; Lyndby et al., 2020). It is thus plausible that this unique cellular organization limits the expulsion of the symbionts and explains the observed high occurrence of *in hospite* symbiont degradation. *In hospite* degradation may be relatively slower than expulsion, which combined with the body shrinkage of the animal may contribute to a delay of visible bleaching in *Cassiopea*.

### An autophagic immune response of *Cassiopea* to heat stress?

The *in hospite* degradation of algal symbionts was accompanied by an overall increase in the density of host nuclei in the mesoglea that was most pronounced in the vicinity of algal symbionts (**Figure 4G**). While this increase in host nuclei density in the mesoglea may be partially ascribed to host body shrinkage, light microscopy and SEM imaging showed that a high number of these nuclei in the oral mesoglea were part of host amoebocytes clusters surrounding the damaged symbiont cells (**Figure 4D, F, S5**). In Anthozoa, amoebocytes have previously been described as effector cells in the immune response, which migrate to sites of injury or infection and can phagocytose foreign, malfunctioning, and damaged cells (Fankboner, 1971; Parisi et al., 2020; Snyder et al., 2021), also in the context of heat stress (Mydlarz et al., 2008; Olano & Bigger, 2000; Palmer et al., 2011; Vargas-Ángel et al., 2007). Hence, we propose that the observed clusters of host amoebocytes in the heat-treated mesoglea can be considered as part of an autophagic *Cassiopea* immune response to the presence of damaged cells in the mesoglea (Downs et al., 2009). We hypothesize that the presence of damaged symbiont cells (and potentially the damaged amoebocytes hosting them) in the mesoglea may attract amoebocytes free of symbionts, to engulf and phagocytose the damaged cells.

At this point, the mechanisms underlying the *in hospite* degradation of algal symbionts clearly require further study. On the one hand, the immune response could be a direct consequence of algal symbionts damaged by heat stress (necrosis). On the other hand, enhanced production of reactive oxygen species (Lesser, 1997; Ventura et al., 2018), reduced nutrient translocation (Rädecker et al., 2021), and parasitic behavior of symbionts (Baker et al., 2018) may provide cues for amoebocyte cells to degrade symbiont cells or initiate host cell apoptosis.

In the sea anemone Aiptasia and symbiotic octocorals, necrosis, and apoptosis seem to co-occur in the heat stress response (Dunn et al., 2002, 2004, 2007; Paxton et al., 2013; Sammarco & Strychar, 2013). Further histological and immunological studies to evaluate amoebocyte mobility and phagocytic activity, as well as the importance of apoptotic or necrotic pathways in symbiosis regulation will be vital to decipher the mechanisms underlying the breakdown of the *Cassiopea* holobiont during heat stress.

### Ecological relevance

While host shrinkage and *in hospite* symbiont degradation may have compensated for and delayed a visible bleaching response, the prolonged exposure to severe heat stress of 34 ºC exceeded the thermal tolerance of the *Cassiopea* holobiont in the present study. While thermal thresholds are species and context dependent, the thermal upper limit measured in this study remains consistent with previous reports documenting limits at 34°C or above (Béziat & Kunzmann, 2022; Klein et al., 2019; McGill & Pomory, 2008). Thus, while *Cassiopea* displayed signs of susceptibility to acute heat stress, their thermal tolerance likely exceeds that of most scleractinian corals (W. Fitt et al., 2001; Klein et al., 2019). By highlighting the importance of the host’s nutritional status during heat stress, our study suggests that the high trophic plasticity of *Cassiopea* may represent an effective way to mitigate host starvation in case of heat stress. In addition, the substantial mesoglea of *Cassiopea* may provide an additional intrinsic energy reservoir that can support the increased metabolic requirements of the host over extended periods with anomalously high water temperatures. Together, its high heterotrophic capacity and large energy reservoir in the mesoglea may confer to *Cassiopea* the ability to thrive in changing, warming and anthropogenically affected environments.

## Conclusion

While the cellular-level events leading to the breakdown of the symbiosis and the associated bleaching phenotype of heat-stressed *Cassiopea* may differ somewhat from other photosymbiotic cnidarians, our results suggest that there are also important similarities with processes described in scleractinian corals and sea anemones (Cziesielski et al., 2022; Rädecker et al., 2021). In particular, during heat stress, energy and carbon limitation of the host rapidly shifts the host metabolism to a net catabolic state in which the relative contribution of algal photosynthates to host nutrition is greatly reduced. This thus reinforces the notion that *Cassiopea* represents a highly relevant laboratory model organism to study the metabolic response of photosymbiotic cnidarians facing heat stress. In addition, our study suggests that the unique anatomical features of *Cassiopea* may represent a clear adaptive benefit, contributing to its ability to tolerate and spread in rapidly changing and extreme environments.

Taken together, the observed ‘invisible’ bleaching phenomenon and relative heat tolerance of *Cassiopea* likely are a result of its unique anatomy and cellular organization. At the same time, this study shows that host bioenergetic status represents a critical parameter shaping the heat stress response among symbiotic cnidarians.

## Acknowledgements

The authors would like to thank N.H. Lyndby for help with the maintenance of the *Cassiopea* culture tank, K. Vernez for the ammonium measurements of seawater samples, J. Daraspe and D. De Bellis for their advice on sample preparation for EM, and C. Göpfert for the advice on histology. Histological sample preparation was performed with the help of the EPFL Histology Core Facility. GT, NR, GBP and AM were supported by the Swiss National Science Foundation grants 200021_179092 and 205321_212614. CP was supported by the Junior Professorship Grant ‘*A connected underwater world’* awarded by the French National Research Agency and an associated start-up grant by the French National Centre for Scientific Research (CNRS).

## Conflict of Interest Statement

The authors have no conflict of interest.

## Authors contributions

GT, NR, CP, GBP, AM conceived the experiment. GT, NR, CP performed the experiments. GT, NR, CP, SE, CG, CMO, JS acquired and analyzed the data. GT wrote the first draft of the manuscript. All authors contributed to reviewing and revising the manuscript.

## Supplementary data

Figure S1 to S5 and Table S1.

## Data availability

All raw data associated with this study have been deposited in the zenodo.org repository: https://doi.org/10.5281/zenodo.8020430.

## Notes

https://doi.org/10.5281/zenodo.8020430

### Competing Interest Statement

The authors have declared no competing interest.

