## Supplementary material for "Host starvation and *in hospite* degradation of algal symbionts shape the heat stress response of the *Cassiopea*-Symbiodiniaceae symbiosis"

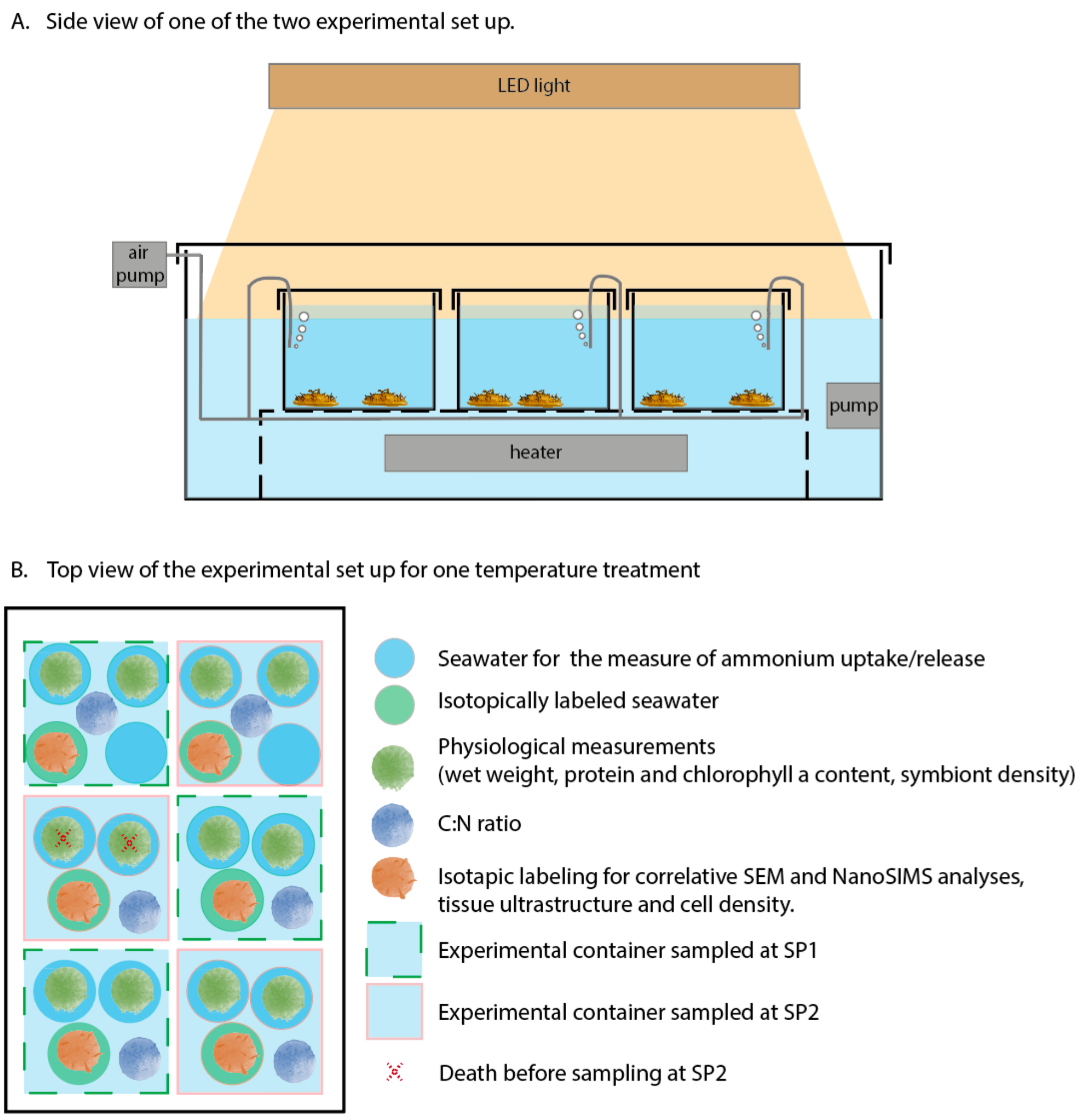


***<u>Figure S1:</u> Schematic presentation of the experimental setup viewed from the side (A) and from above with sampling design information (B).***

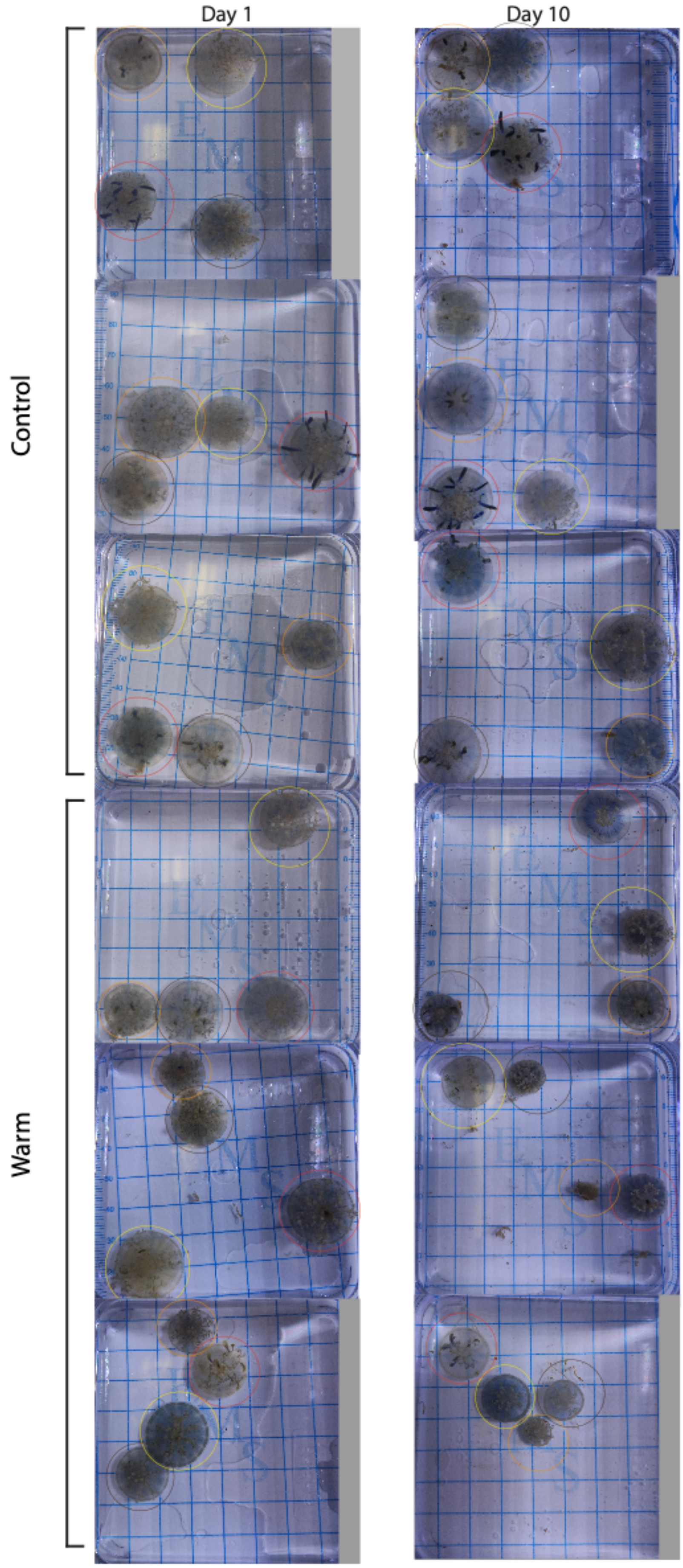


***Figure S2: Photos of the medusae collected at SP2 on day 1 and 10*** *of the experiment in the control treatment, and heat stress treatment. Circles of the same color within each line indicate the same medusa.*

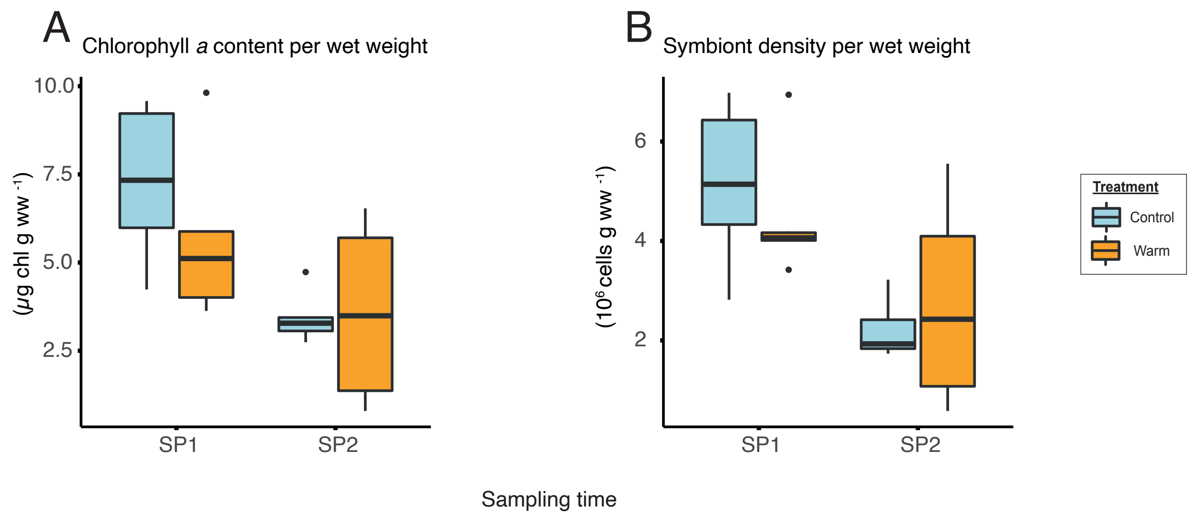


***<u>Figure S3</u> : Impact of heat stress on chlorophyll a content (A) and algal symbiont density (B) per wet weight***

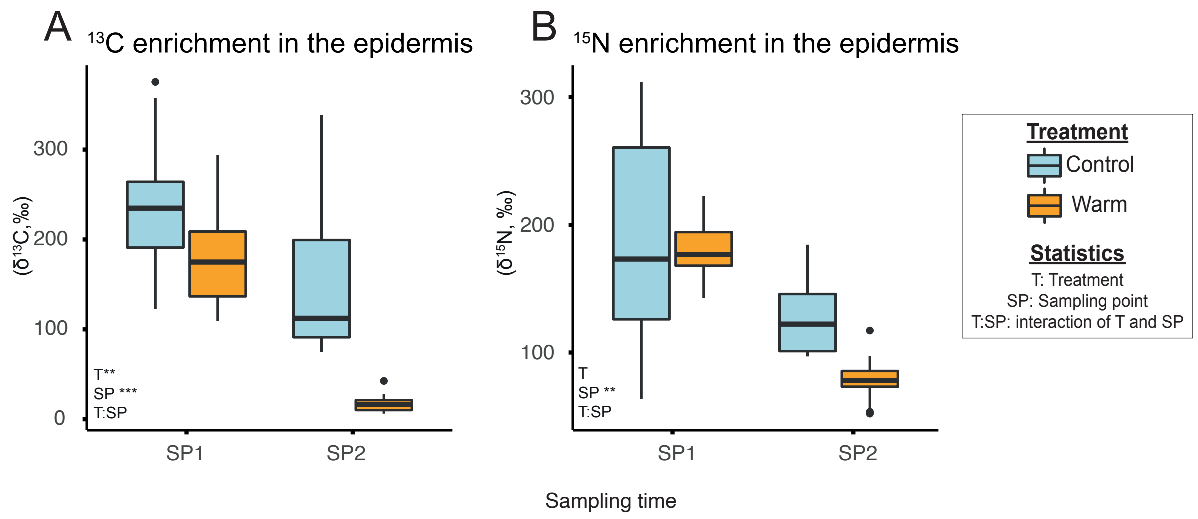


***<u>Figure S4</u>*: *Temperature effects on the enrichment from assimilation of isotopically labeled bicarbonate and ammonium in the epidermis of heat-stressed medusae.*** *$^{13}$C enrichment in from assimilation of $^{13}$C-bicarbonate ($H^{13}CO_3^-$) via photosynthesis and $^{15}$N enrichment induced by the assimilation of $^{15}$N-Ammonium ($NH_4^+$) into algal symbiont cells and host amoebocytes. Asterisks indicate significant differences between treatments (* $p < 0.050$, ** $p < 0.010$, *** $p < 0.001$).*

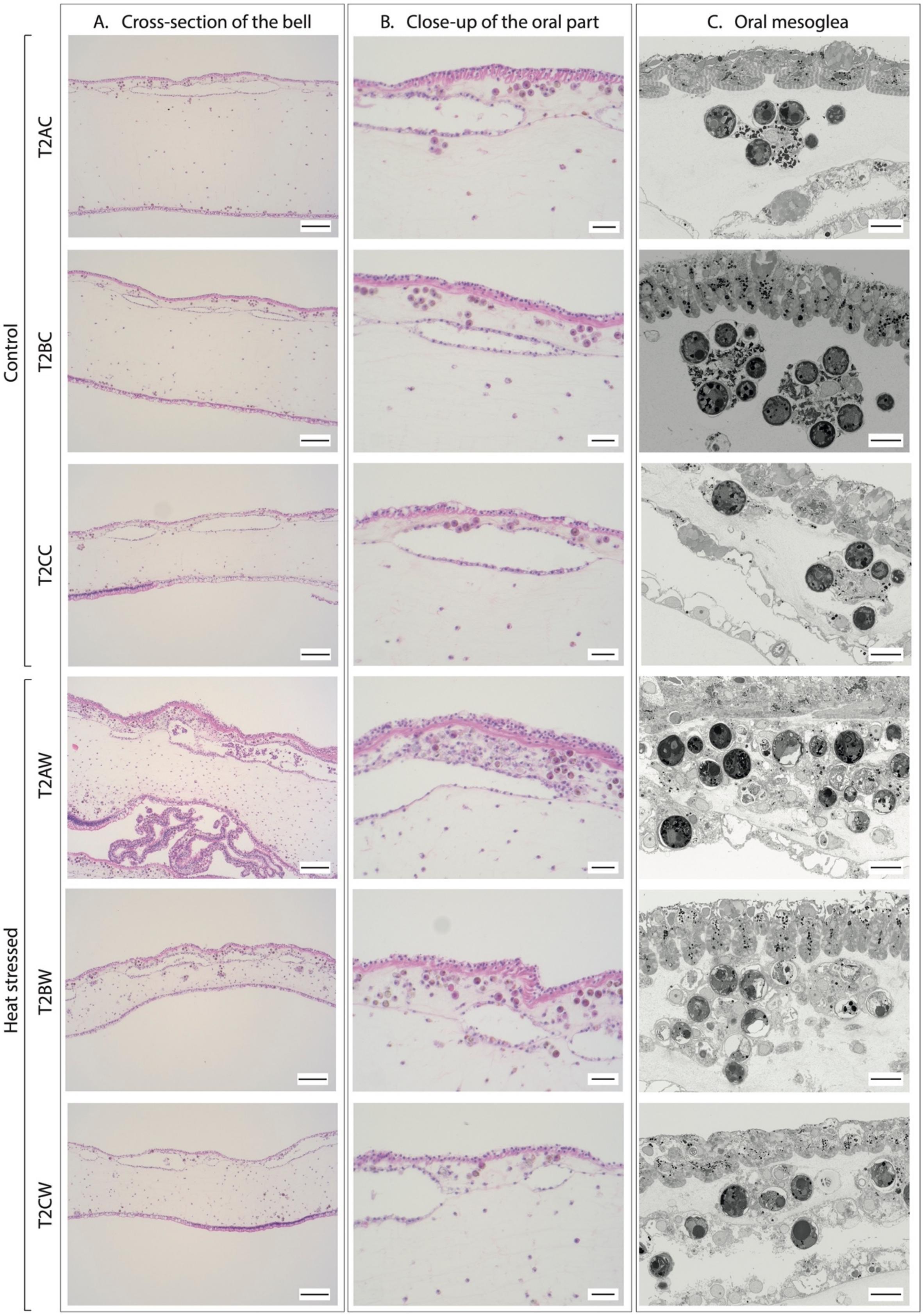


***<u>Figure S5:</u> Tissue and cellular ultrastructure of control and heat-stressed medusae bells at SP2*** *imaged by light microscopy with H&E staining (A,B) and SEM (C). Each row corresponds to one biological replicate. The annotations on the left-*

*hand side of each row correspond to the unique sample numbers of individual replicate medusae. Scale bar A: 100μm, B: 25μm, C: 10μm.*

***Table S1:*** *Number of NanoSIMS images per biological replicate and the associated number of regions of interest (ROIs) per* Cassiopea *compartment defined for the isotopic enrichment analyses.*

| | Biological replicates | Number of images | ROIs Symbiont | ROIs Amoebocytes | ROIs Epidermis |
|---|---|---|---|---|---|
| SP1 Control | A | 7 | 36 | 8 | 5 |
| | B | 6 | 25 | 6 | 6 |
| | C | 6 | 29 | 7 | 5 |
| SP1 Warm | A | 5 | 31 | 6 | 5 |
| | B | 5 | 23 | 5 | 5 |
| | C | 6 | 25 | 6 | 5 |
| SP2 Control | A | 5 | 28 | 5 | 5 |
| | B | 6 | 33 | 7 | 4 |
| | C | 6 | 12 | 6 | 6 |
| SP2 Warm | A | 9 | 23 | 7 | 9 |
| | B | 8 | 34 | 7 | 5 |
| | C | 3 | 3 | 2 | 1 |
| Unlabeled | | 26 | 91 | 24 | 23 |
